# Sustained Humoral Activation through self-amplifying mRNA Vaccination Enhances Longitudinal Antibody Function in a Phase III Trial

**DOI:** 10.1101/2025.11.25.690547

**Authors:** Kate S. Levine, Ross Blanc, Qixin Wang, Hadar Malca, Rose Sekulovich, Hongfan Jin, Sophie Liu, Carole Verhoeven, Brian Sullivan, Igor Smolenov, Ryan P. McNamara

## Abstract

Sustained and functional antibody responses to respiratory pathogens through vaccination is critical for global public health. The development and deployment of mRNA vaccines during the coronavirus disease 2019 (COVID19) pandemic was a landmark achievement in modern medicine and ushered in a new age of vaccine innovation. The mRNA-based vaccines elicited strong antibody responses, both neutralizing and extra-neutralizing, against the viral Spike protein. The antibody levels waned with time since vaccination, and that coupled with the antigenic drift of the virus prompted updates to the mRNA vaccine composition and evaluation of other mRNA modalities. Self-amplifying mRNA (sa-mRNA) vaccines such as ARCT-154 can prolong antigen production and durability of humoral immune response post-immunization, and can thus be administered at a lower dose. How this translates into the overall humoral architecture compared to that shaped by conventional mRNA vaccinations, however, is unclear. Here, we analyze serum-based antibody responses from a recent Phase III trial comparing humoral responses elicited by ARCT-154 and mRNA BNT162B2. All participants had received three doses of mRNA COVID-19 vaccines and were randomized to receive a booster dose of ARCT-154 or BNT162B2. Primary outcomes were to quantify waning responses against ancestral/wild type SARS-CoV-2 Spike (WT Spike) and a panel of antigenically SARS-CoV-2 variants Spikes. Through a systems serology approach, we identified that the sa-mRNA vaccine ARCT-154 elicited a unique antibody response compared to BNT162B2 defined by a sustained, activating profile to the vaccine-encoded Spike protein and a broad spectrum of drifted Spikes. Notably, potently activating FcγRIIIA-binding antibodies showed a sustained stimulation in the ARCT-154-treatment arm, and this translated to an enhanced natural killer (NK) cell activation. The NK-activation through ARCT-154 was present for both target WT Spike and the antigenically drifted BA.5 Spike, which was the predominant form of SARS-CoV-2 during the observation period. Our results support a model whereby prolonged antigen expression and presentation moves immune profiles towards activating phenotypes with broad antigenic coverage.

## Introduction

The rapid development and deployment of effective mRNA vaccines against severe acute respiratory syndrome coronavirus 2 (SARS-CoV-2) were instrumental in curbing the coronavirus disease 2019 (COVID-19) pandemic. The mRNA vaccines encode for the SARS-CoV-2 spike protein (S-protein) to induce cellular and humoral immune responses. The vaccines elicited robust antibody titers, including neutralizing antibodies, to Spike that waned with time since immunization (1–7). As the virus further adapted to its new human host, variants of concern (VOC) emerged that showed degrees of neutralizing antibody escape. This was most evident during the emergence of the Omicron lineage of SARS-CoV-2 (8–15). To that end, vaccine boosters and updated vaccine compositions encoding for new emergent SARS-CoV-2 variants were utilized to restore waned immunity and expand breadth (16–23).

Self-amplifying mRNA (sa-mRNA) vaccines encode a target immunogen along with an RNA-dependent RNA polymerase to allow for longer periods of antigen production (24). This is commonly coupled with a decrease in the total mRNA dose required to induce immunity, thereby improving the deployment of mRNA vaccine technology. One such candidate, the sa-mRNA ARCT-154, was previously analyzed for safety, efficacy, and immunogenicity as a primary series and a booster dose. In the pivotal primary vaccination study, ARCT-154 produced higher and more durable neutralizing antibody response compared to the adenovirus vector-based vaccine, which translated into higher efficacy against symptomatic COVID-19 disease (25). Separately, recipients who had previous mRNA immunizations were boosted with ARCT-154 and had higher antibody titers to the target Spike (WT Spike) and to the BA.4/5 Spike variant 28 days after immunization than the conventional mRNA comparator (26), and this trend was sustained for up to 12 months post-boost (27, 28). Other preclinical-stage sa-mRNA vaccine candidates have also shown an ability to elicit immune responses at lower doses compared to conventional mRNA delivery platforms, further supporting the lower-cost, equitable distribution of the vaccines (24).

While neutralizing antibodies are frequently assayed as a primary correlate of protection, effector functions of antibodies have been shown to play critical roles in protection against respiratory pathogens such as SARS-CoV-2 and influenza virus (7, 29–35). It has been shown that mRNA vaccines can elicit and boost both neutralizing and effector-driven antibodies, thereby providing a layered humoral protection strategy (36–41). These effector functions are driven by engagement of the antibody Fc domain with Fc-receptors (FcR) on the surface of innate immune cells such as monocytes, neutrophils, and natural killer cells. Studies have shown that waned neutralizing and extra-neutralizing functions can be recalled and expanded through mRNA vaccine boosters against SARS-CoV-2 (39, 42–45).

In this study, we interrogate the antibody repertoire of the Phase III booster trial of recipients of the sa-mRNA ARCT-154 (also known as Kostaive) or mRNA BNT162B2 (also known as Comirnaty). Participants had received three doses of licensed mRNA COVID-19 vaccines before enrollment, and were randomized into two treatment groups. Serum collections were performed at day 1 (baseline, day of booster), day 29, day 91, day 181, and day 361 after the booster. The main analyses were performed during the waning period (days 29-361) to quantify durability of antibody responses to the direct target (defined as WT Spike) and target-related Spikes (defined as Delta, BA.2, BA.5, XBB.1.5, and KP.3). The direct target and target-related humoral responses were compared to non-target control antigens such as influenza hemagglutinin (high-exposure rate antigen), human cytomegalovirus glycoprotein (high exposure rate antigen), and Ebolavirus glycoprotein (low/negative exposure rate antigen). Our analysis revealed distinct humoral immune responses between the two vaccine groups, contributing to differences in the magnitude, durability, and breadth of effector and neutralizing antibody responses. Many of these distinctions persisted through the entirety of the observation period, up to day 361 post-boost. Notably, ARCT-154 recipients exhibited a unique humoral stimulation and waning rate compared to BNT162b2 recipients, both to the direct target (WT Spike) and target-related (Delta and Omicron Spikes). These findings underscore the divergent humoral immune profiles elicited by sa-mRNA and mRNA booster regimens and provide insight into how sustained humoral stimulation translates to immunity across SARS-CoV-2 variants.

## Results

### Humoral Responses to mRNA and sa-mRNA Vaccines are Distinct for up to 180 Days Post-Immunization

A phase III non-inferiority trial comparing responses to COVID-19 boosters was performed (see Methods), and specimens were analyzed through systems serology. All participants had received a 2-dose primary vaccination series with mRNA COVID-19 vaccines, and a booster dose of BNT162B2 vaccine at least three months before enrollment. Participants were randomized into two groups: those receiving a booster dose of BNT162b2 and those receiving a booster dose of the self-amplifying mRNA ARCT-154 (25–28). Blood draws were taken at day 1 (baseline, day of booster dose), day 29 post-boost, day 91 post-boost, day 181 post-boost, and day 361 post-boost.

Evaluation of Ig isotypes (IgG, IgA, IgM), subclasses (IgG1-4), and low-affinity Fc gamma receptor binding (FcγR) antibodies to on-target (WT Spike), target-related (Delta Spike, BA.2 Spike, BA.5 Spike, XBB.1.5 Spike, and KP.3 Spike), and non-target (influenza H1N1 A/Wisconsin/22 hemagglutinin (HA), human cytomegalovirus (HCMV) Glycoprotein B (GB), and Ebolavirus Glycoprotein (GP)) controls was assessed (**Figure 1**). Influenza H1N1 HA and HCMV GB served as consistency non-target controls, which should be present in the vast majority of participants, but not responsive to the boosters, while Ebola GP served as a non-target negative control. Additionally, antibody-dependent phagocytosis by monocytes (ADCP) and antibody-dependent natural killer cell activation (ADNKA) to target (WT), target-related (BA.5), and non-target (Ebola GP) were assessed. During the observation period of this study, BA.5 was the predominant form of SARS-CoV-2.

**Figure 1.**
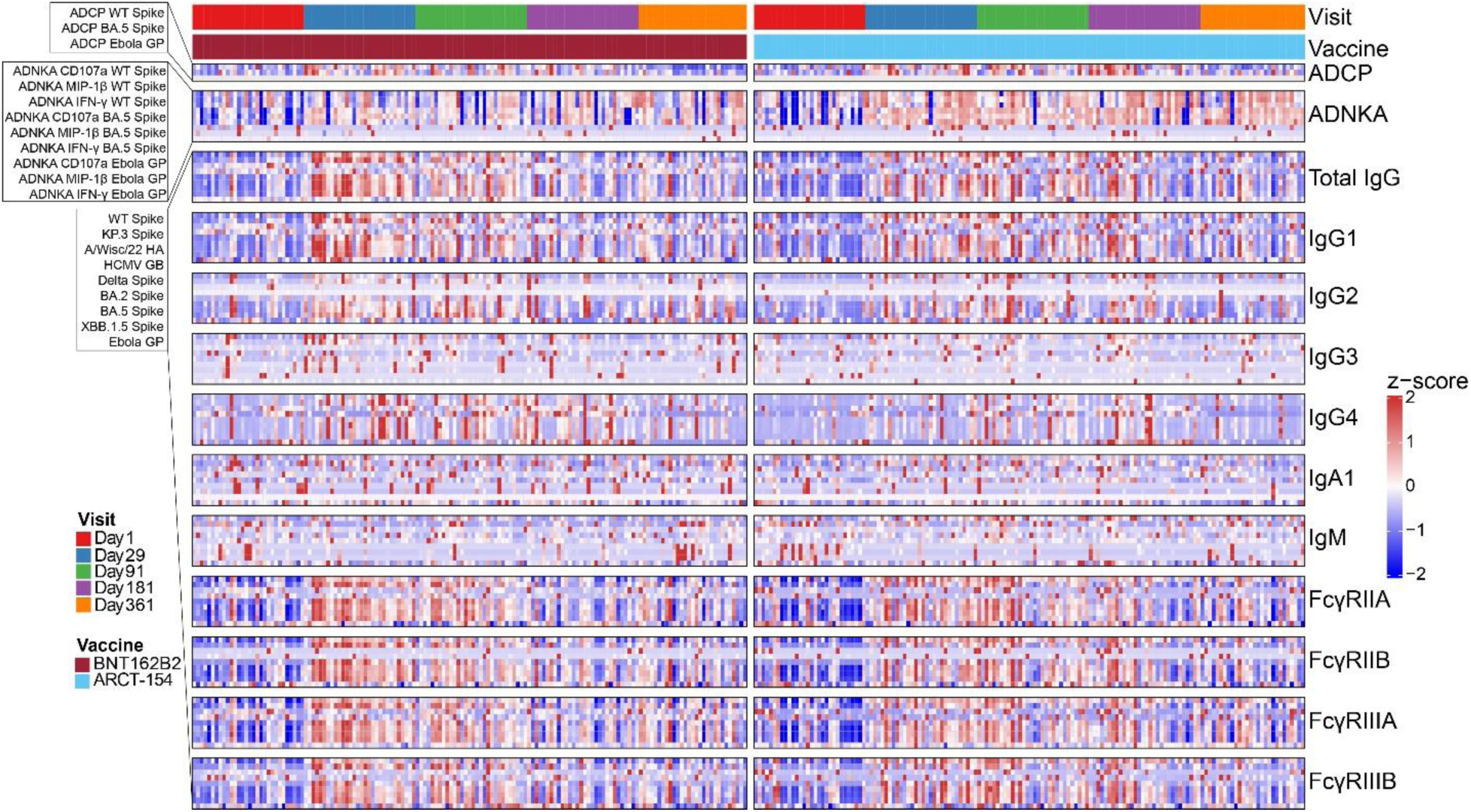
Systems serology analysis of the ARCT-154-J01 boosting trial. Participants who had received two doses of licensed mRNA COVID-19 vaccines, and a third dose of BNT162B2 at least three months before the study recruitment were randomly assigned to two treatment arms: receiving a booster dose of BNT162B2 (red block, left) or ARCT-154 (blue block, right). Blood draws were taken at baseline (day 1), and at days 29, 91, 181, and 361 post-boost. Shown are effector function and binding profiles in a scaled heatmap. Each column represents one participant within the designated treatment group at the indicated time, and each row is the humoral feature. On the left are the ordering for analytes for antibody-dependent cellular phagocytosis by monocytes (ADCP, top row), antibody-dependent natural killer cell activation (ADNKA, second row), and antibody binding (remaining rows). All outputs were z-scored across rows, and the scaling legend is shown on the right. Values in gray indicate that the readout did not pass quality control, meaning that replicate standard deviations exceeded 50% of the mean, or that values were not above our lower limit of quantitation (see Methods).

Overall antibody binding and functional architecture revealed stimulation at 29 days post-vaccination for both treatment groups (BNT162B2 in red, ARCT-154 in blue) with responses waning thereafter. This stimulation and waning showed antibody isotype, subclass, and FcγR specificity, with not all features showing similar trends, as expected.

To identify if the overall antibody profile could be separated between the two boosted groups, we performed a partial least squares discriminant analysis (PLSDA) model of antibody binding. A least absolute shrinkage and selection operator (LASSO) regularization was used to down-select the number of antibody features for the PLSDA fit (46, 47) (see Methods). This was done for each draw during the observation period (**Figures 2A-D**, Supplementary Figure 1). No model could be built distinguishing recipients of BNT162B2 and ARCT-154 for day 1, indicating that baseline humoral profiles were not significant between groups. However, multivariate clustering through this method could statistically separate out treatment groups (BNT162B2 in red, ARCT-154 in blue) at day 29-, 91-, and 181-post-vaccination (for each, *p* < 0.04 between the model and random features, *p* < 0.02 between the model and permuted labels). A model could not be built, however, at 361 days post-boost (*p* = 0.1 between the model and random features, *p* = 0.07 between the model and permuted labels) (**Figures 2E – 2H**).

**Figure 2.**
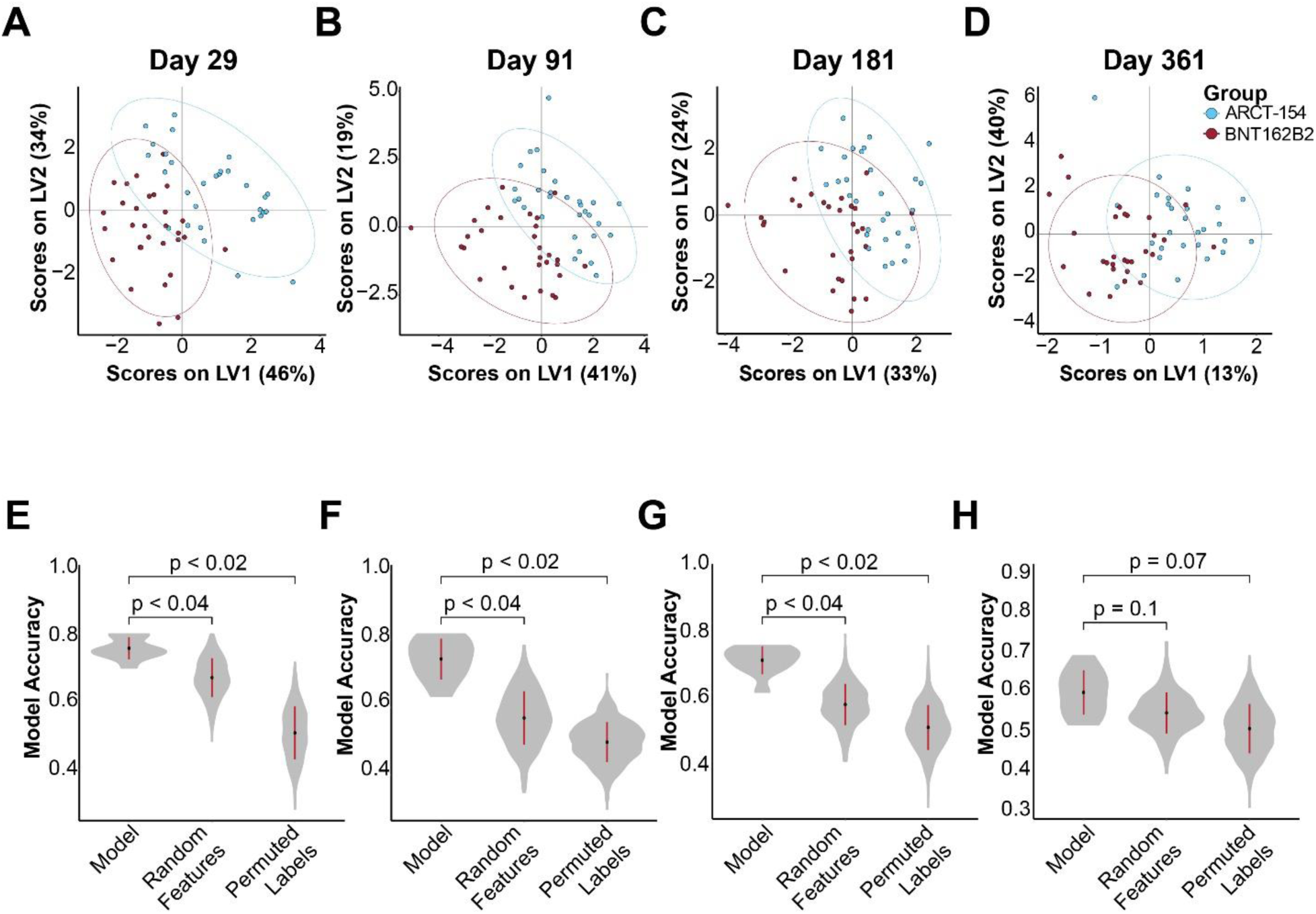
Recipients of the mRNA vaccine BNT162B2 and the sa-mRNA vaccine ARCT-154 generate distinct humoral profiles longitudinally. (A) Probable least squares discriminant analysis (PLSDA) was used to identify clustering based on treatment groups on day 29. In red dots is the composite humoral profile (binding only) of recipients of BNT162B2, and in blue dots are recipients of ARCT-154. Multivariate separation is shown on latent variable (LV) axes 1 and 2. Ellipses indicate the 95% confidence intervals of the multivariate plotting of the group. (B) Same as A, but for antibody binding profiles at day 91. (C) Same as A, but for antibody binding profiles at day 181. (D) Same as A, but for antibody binding profiles at day 361. (E) Validation of PLSDA and LASSO-selected features for A. The LASSO-selected features describing separation at the multivariate level were used to quantify model accuracy (left column). The model was then compared to one of random features (middle column) and permutated labels (right column). P-values are shown above the comparisons. (F) Same as E, but for model performance of the LASSO PLSDA at day 91. (G) Same as E, but for model performance of the LASSO PLSDA at day 181. (H) Same as E, but for model performance of the LASSO PLSDA at day 361.

The LASSO-selected features contributing to separation at the multivariate scale were plotted along latent variable axes 1 and 2 (LV1 and LV2, respectively). FcγR-binding antibody responses to target WT Spike were selected in the ARCT-154 recipients at each timepoint, while antibody binding features to Omicron lineages were selected in the BNT162B2 arm (Supplementary Figure 1). These results suggested that the profiles were moving at distinctly different rates after the boost and before the end of the observation period. Also, even though ARCT-154 and BNT162B2 were directed to WT/D614G Spike, there were distinctions in the humoral architecture to it and to target-related Spikes that could influence function.

### sa-mRNA Vaccination Yields a Sustained Stimulation to On- and Related-Targets

It was previously reported that recipients of the sa-mRNA vaccine ARCT-154 had a more durable neutralizing antibody phenotype compared to other mRNA-based vaccines (27, 28). Our data supported that rates of change to target and target-related Spikes between the two treatment arms may be driving separation at the multivariate level. We thus took the first derivative (*d*) of the collection periods to quantify how rates of change of antibody levels may differ between sa-mRNA and mRNA booster recipients. We were particularly interested in how activating FcγR-binding antibodies moved with time, given the previously reported difference in neutralization between an mRNA and a sa-mRNA vaccination platform (26–28). Our primary analysis was thus post-peak responses (after day 29) to the target (WT Spike) and target-related (Delta, BA.2, BA.5, XBB.1.5, and KP.3) Spikes during this window.

Total IgG levels were first analyzed for rates of change between the collections. The ARCT-154 treatment arm exhibited a sustained total IgG stimulation to target WT Spike in the time interval of day 29 to day 91 (median *d* > 0), whereas recipients of BNT162B2 displayed a sharp contraction in total IgG to the target antigen (median *d* < 0). This was statistically significant between the two treatment arms (false-discovery rate, or FDR, corrected p-value < 0.05).

Subsequent time intervals showed a median *d* < 0 for both treatment arms, indicating contracting total IgG levels. The *d* Total IgG at day intervals 91-181 and 181-361 were not statistically significant (**Figure 3A**). Interestingly, this same trend was observed for each target-related Spike (**Figure 3B-F**). For these antigenically drifted Spike variants, the sa-mRNA ARCT-154 recipients showed a median *d* Total IgG ≥ 0 in the day 29-91 time interval, indicating that median waning was not occurring for the group until after day 91. This was in contrast to BNT162B2, which showed a *d* Total IgG < 0 at the day 29-91 time interval, and either further contracted or showed a flat rate of Total IgG levels thereafter. There were no changes in *d* Total IgG for HA A/Wisconsin/22 or HCMV GB, indicating that non-target, high exposure antigen responses were neither stimulated nor contracted (**Figure 3G, H**). Lastly, no changes were observed for non-target, negative control Ebola GP (**Figure 3I**).

**Figure 3.**
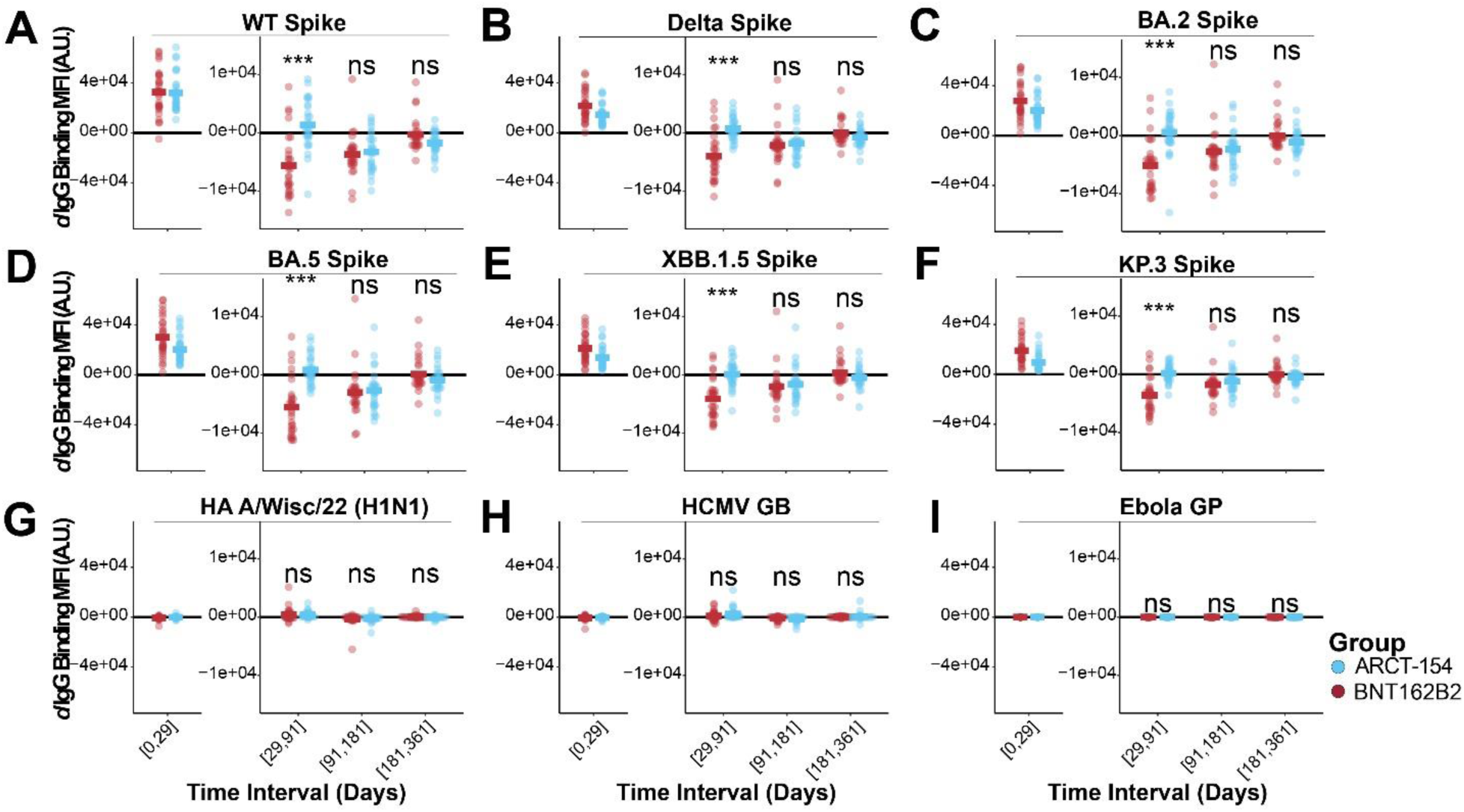
IgG levels to target and target-related antigens have different rates of decay based on boosting with an mRNA- or an sa-mRNA-vaccine. (A) Derivatives (rate of change, or *d*) were quantified for the time intervals shown on the x-axis for the two treatment arms (BNT162B2 in red, ARCT-154 in blue). Each dot represents the *d* of Total IgG to WT Spike at the time interval, and the colored horizontal bar indicates the group mean of *d* Total IgG WT Spike. A *d* < 0 indicates a negative slope, or contraction; a *d* > 0 indicates a positive slope, or expansion; and a *d* = indicates no change in slope, or sustained response. Statistical comparisons between the two groups were done for each time interval using a Wilcoxon Test followed by a false discovery rate (FDR) adjustment. Above each time interval, n.s. indicates not statistically significant after FDR correction (p ≥ 0.05), * indicates p < 0.05 after FDR correction, ** indicates p < 0.01 after FDR correction, and *** indicates p < 0.001 after FDR correction. Acute phase stimulation comparisons were not performed as this was not a primary endpoint analysis (see Methods). (B-F) Same as A, but for variant SARS-CoV-2 Spikes shown at the top of each graph. (G-I) Same as A, but for high-exposure control off-target antigens (HA A/Wisc/22 H1N1 and HCMV GB), and for no-exposure control antigens (Ebola GP).

The same approach was used for IgG subclasses 1-4. A similar trend for IgG1, IgG2, and IgG4 was observed for the sa-mRNA ARCT-154 recipients with sustained antibody levels through the day 29-91 time interval. Only minor IgG3 rates of change were observed, and they were exclusive to Delta and KP.3 Spike in the day 29-91 time interval (Supplementary Figures 2-5). These results indicate that sa-mRNA and mRNA vaccinations elicit differing stimulation and decay dynamics for binding antibodies to target (WT Spike) and target-related (Delta, BA.2, BA.5, XBB.1.5, and KP.3) antigens, but have minimal non-target responses.

Similarly, the derivative plots for the Spike proteins showed discrepancies for activating FcγRIIIA-binding antibodies between the two treatment arms for the target WT Spike (**Figure 4A**). Surprisingly, this *d* FcγRIIIA between the two treatment groups remained statistically significant at the day 91-181 time interval, indicating that the rate of contraction for recipients of ARCT-154 was lower than for BNT162B2 at extended time periods. The two groups both approached *d* FcγRIIIA WT Spike = 0 at the day 181-361 time interval, with BNT162B2 recipients having a higher *d* FcγRIIIA WT Spike at that interval (**Figure 4A**). Sustained FcγRIIIA-binding levels to target-related Spikes also showed statistically significant differences in durability for recipients of the sa-mRNA ARCT-154 at the day 29-91 interval, and a few even showed statistically significant differences at the day 91-181 interval. These rates eventually flatline at days 181-361 (**Figure 4B-F**). There were no changes in *d* FcγRIIIA-binding antibodies observed in our control antigens, which indicates that there were negligible non-target binding fluctuations occurring (**Figure 4G-I**).

**Figure 4.**
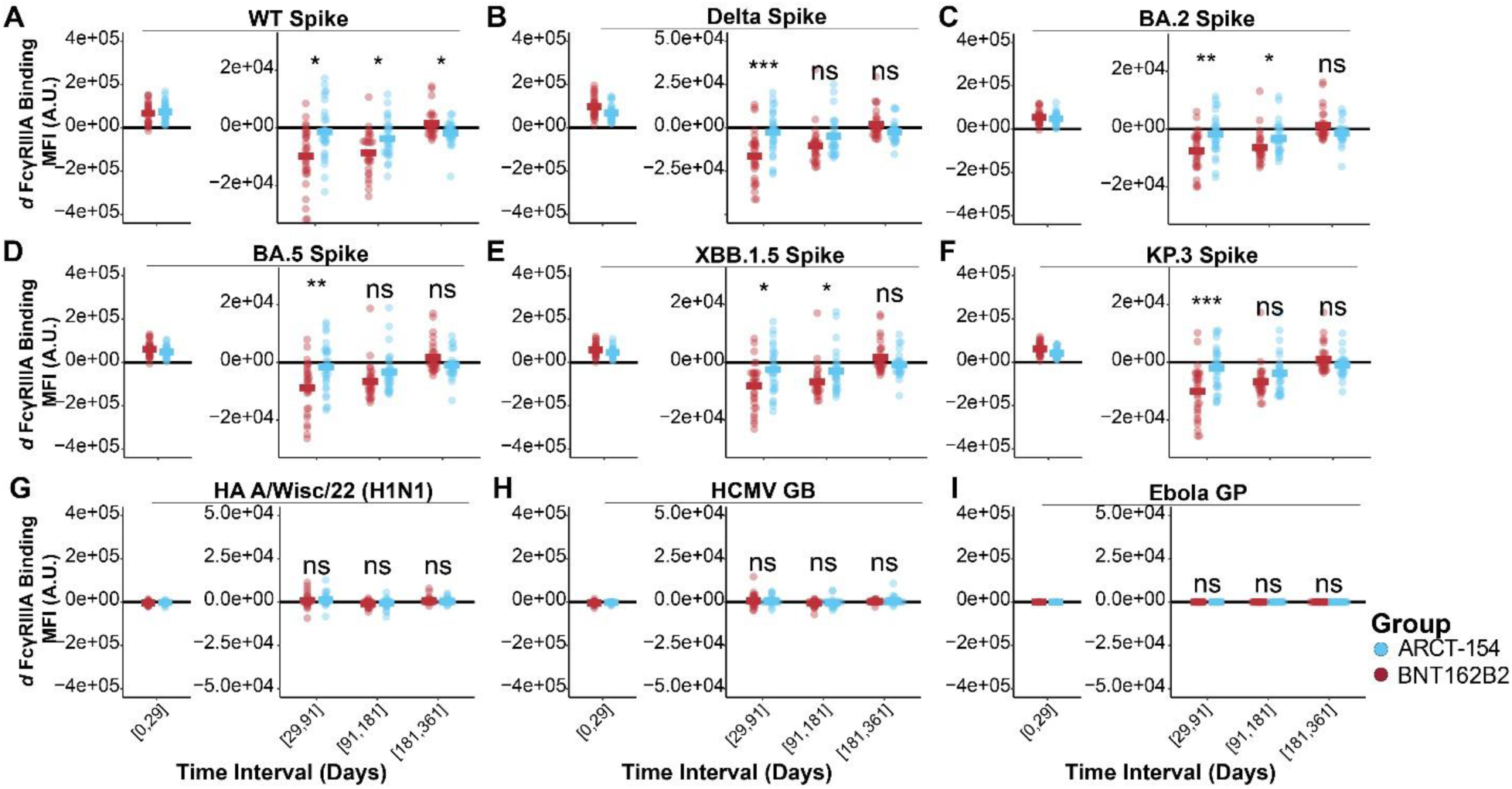
Activating FcγRIIIA binding antibody levels to target and target-related antigens have different rates of decay based on boosting with an mRNA- or an sa-mRNA-vaccine. (A) *d* FcγRIIIA-binding antibody levels were quantified for the time intervals shown on the x-axis for the two treatment arms (BNT162B2 in red, ARCT-154 in blue). Each dot represents the *d* of FcγRIIIA-binding antibodies to WT Spike at the time interval, and the colored horizontal bar indicates the group mean of *d* FcγRIIIA-binding antibodies to WT Spike. A *d* < 0 indicates a negative slope, or contraction; a *d* > 0 indicates a positive slope, or expansion; and a *d* = indicates no change in slope, or sustained response. Statistical comparisons between the two groups were done for each time interval using a Wilcoxon Test followed by a false discovery rate (FDR) adjustment. Above each time interval, n.s. indicates not statistically significant after FDR correction (p ≥ 0.05), * indicates p < 0.05 after FDR correction, ** indicates p < 0.01 after FDR correction, and *** indicates p < 0.001 after FDR correction. Acute phase stimulation comparisons were not performed as this was not a primary endpoint analysis (see Methods). (B-F) Same as A, but for variant SARS-CoV-2 Spikes shown at the top of each graph. (G-I) Same as A, but for high-exposure control antigens (HA A/Wisc/22 H1N1 and HCMV GB) and for no-exposure control antigens (Ebola GP).

Other FcγR-binding antibodies showed differing rates of antibody levels during the observation period. FcγRIIA-binding antibodies showed a trend toward enrichment for ARCT-154 recipients at the day 29-91 interval, with a few Spikes showing statistical significance compared to BNT162B2 recipients, but not nearly to the same degree as FcγRIIIA (Supplementary Figure 6). FcγRIIB showed a statistically significant enhancement in *d* for all Spikes at the day 29-91 time interval for ARCT-154 recipients, but not at any other time (Supplementary Figure 7). Lastly, no differences in *d* FcγRIIIB-binding antibodies were observed between the two treatment groups (Supplementary Figure 8). Importantly, there were no changes in *d* FcγIIA, *d* FcγRIIB, FcγRIIIA, or *d* FcγRIIIB for any of our non-target control antigens. These data indicate that the strongly activating FcγRIIIA was disproportionately sustained in recipients of the sa-mRNA vaccine boosted. This response was specific to on-target and related-target Spikes.

### Antibody Effector Functions Show a Slow Waning Profile after sa-mRNA Boosting

We then profiled the functionality of the humoral response longitudinally over time as Fc-mediated effector functions have been shown to confer protection against SARS-CoV-2 (48–51). Antibody-dependent cellular phagocytosis by primary-derived monocytes (ADCP – see Methods and Supplementary Figure 9A) to target WT Spike and target-related BA.5 Spike had similar initial responses for the two groups. Baseline values were well above our lower limit of detection since the participants all had previous immunity through vaccination. Both groups showed a waning period from days 181 to 361 post-boost. Although trends to elevated ADCP were observed for recipients of ARCT-154, the two groups’ responses were not statistically significant after FDR correction to WT or BA.5 Spike (**Figure 5A**, left and middle). Neither group had any detectable ADCP to our negative exposure control antigen Ebola GP (**Figure 5A**, right).

**Figure 5.**
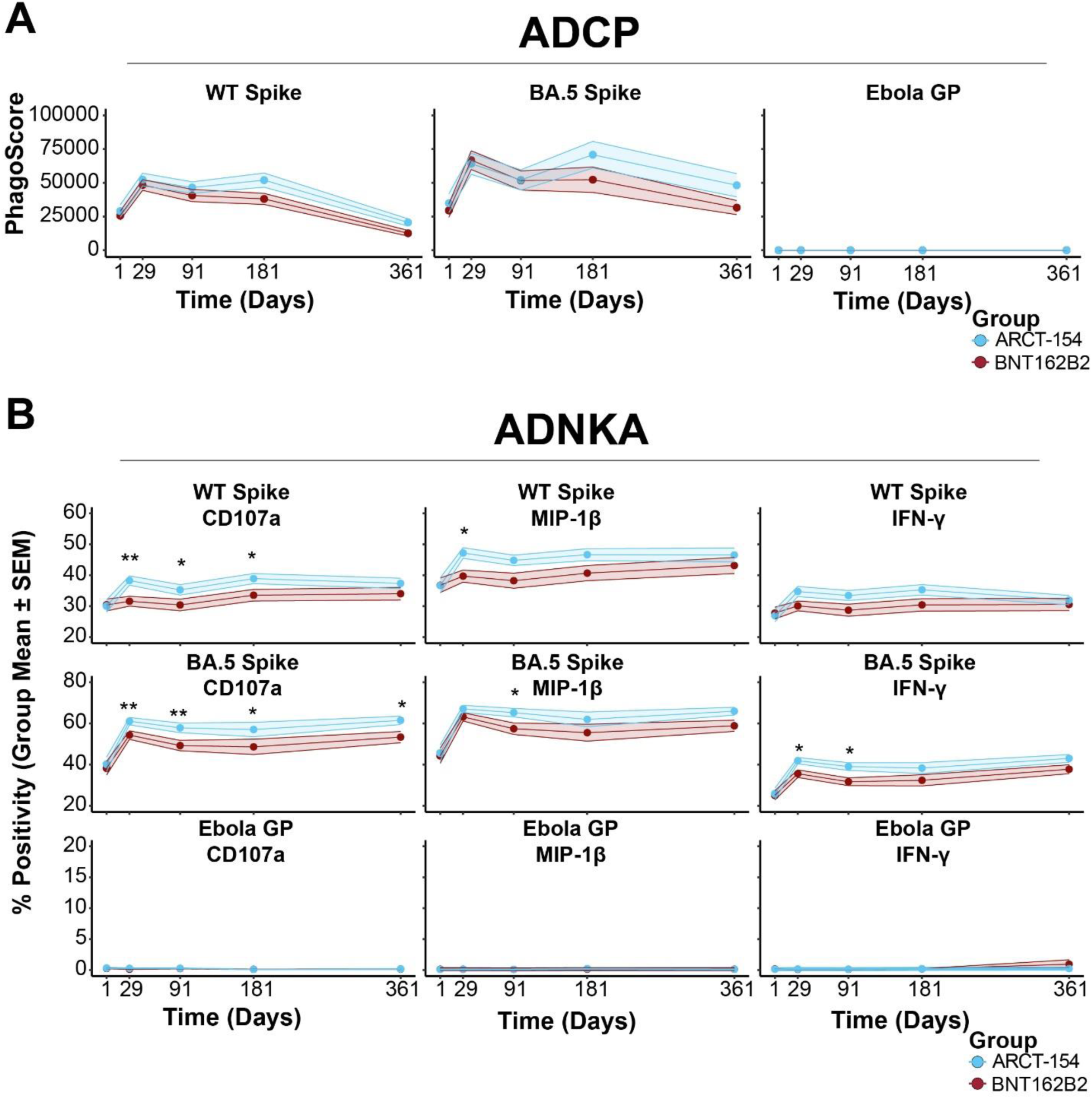
Antibody effector functions to WT and BA.5 Spike are boosted by mRNA and sa-mRNA vaccines, with a sustained phenotype in the latter. (A) Antibody-dependent cellular phagocytosis by monocytes (ADCP) was assayed for the two treatment arms (BNT162B2 in red, ARCT-154 in blue). Shown are group PhagoScore means (y-axis) at each time point (x-axis) along with 95% confidence intervals. ADCP was quantified for target WT Spike (left), target-related BA.5 Spike (middle), and non-target Ebola GP (right). (B) Antibody-dependent natural killer cell activation (ADNKA) was assayed for the two treatment arms (BNT162B2 in red, ARCT-154 in blue) through surface expression of CD107a, macrophage inflammatory protein 1 beta (MIP-1β) expression, and interferon gamma (IFN-γ) expression. Shown are the group percent positivity means (y-axis) at each time point (x-axis) along with 95% confidence intervals. ADNKA was quantified through the three outputs for target WT Spike (top row), target-related BA.5 Spike (middle row), and Ebola GP (bottom row). Above each time point, statistical differences between treatment arms were noted. * indicates p < 0.05 after FDR correction, ** indicates p < 0.01 after FDR correction, and *** indicates p < 0.001 after FDR correction. Shown are the group means (dots) at each time point along with 95% confidence intervals (shaded region).

Cellular cytotoxic activity was quantified through an antibody-dependent natural killer cell activation (ADNKA) assay in which percent positivity of degranulation marker CD107a, macrophage inflammatory protein 1 beta (MIP-1β), and pro-inflammatory cytokine interferon gamma (IFN-γ) were measured (Supplementary Figure 9B). ADNKA activity to the target WT Spike for ARCT-154 recipients showed a significant stimulation at day 29 compared to BNT162B2 recipients for CD107a and MIP-1β. A persistence of the CD107a marker (cellular degranulation) was noted until day 181 for the ARCT-154 treatment arm. A similar trend, albeit not statistically significant after FDR correction, was observed for IFN-γ production in response to target WT Spike exposure. (**Figure 5B**, top row). It is important to note that both treatment arms had high pre-existing immunity to the target WT Spike.

We next quantified ADNKA activity to the target-related BA.5 Spike. Here, both treatment arms exhibited a strong acute response for all three ADNKA outputs, indicating that the breadth of responses was increasing, consistent with previously reported observations (39, 44, 52). Similar to other assays, ADNKA responses were relatively sustained in the ARCT-154 treatment arm.

For CD107a expression, recipients of ARCT-154 exhibited a significantly higher response to the target-related BA.5 Spike compared to the BNT162B2 recipients for the entire observation period. All three ADNKA markers showed enrichment for ARCT-154 recipients at day 91 post-boost (**Figure 5B**, middle row). Lastly, ADNKA responses were quantified to an off-target antigen, Ebolavirus GP. Here, neither treatment arm had detectable baseline ADNKA activity, nor were they stimulated by vaccination at any timepoint (**Figure 5B**, bottom row).

Taken together, our functional assays closely tracked with binding antibody profiles for recipients of a BNT162B2 or ARCT-154 booster after receiving three doses of an mRNA vaccine. Notably, ADNKA remained elevated for recipients of the sa-mRNA ARCT-154 booster, which supported the slower decay dynamics of FcγRIIIA-binding antibodies observed. Also, antibody effector functions were specific to SARS-CoV-2 for both treatment arms.

## Discussion

Eliciting durable and functional antibody responses is a point of focus for next-generation vaccination strategies and platforms. The development and deployment of mRNA-based vaccines against COVID-19 have ushered in a new era of vaccinations, and understanding how the immune responses shaped by the vaccine can translate to protection. The mRNA-based vaccines showed strong protection against SARS-CoV-2 in clinical trials and in real-world data (2, 3, 5, 6, 53–57). As the virus adapted itself to its new host, variants swept through the population that exhibited increased transmissibility and decreased recognition by neutralizing antibodies, possibly through intrapatient evolution (9, 10, 53, 58, 59). The evolving antigenic targets of the Spike glycoprotein, coupled with the waning of antibody responses acquired through vaccination or infection, have prompted updates to mRNA-vaccine technology. This has come in the form of updating the Spike open reading frame encoded in the mRNA (21–23, 60), as well as encoding an RNA-dependent RNA polymerase as a separate open reading frame to self-amplify the mRNA and increase production of the Spike (24–28).

Previous work showed that a self-amplifying mRNA (sa-mRNA) vaccine could enhance the durability of neutralizing antibodies (26–28), which were a correlate of protection against ancestral SARS-CoV-2 (7). Antibody effector functions have also been shown to exert protection, particularly against antigenically diverged Spikes like the Omicron lineage (50, 61), and are recalled upon infection (42, 49). The goal of this study was to interrogate whether a sa-mRNA vaccine could elicit a sustained and activating antibody response within a population where responses could be recalled.

We found that the sa-mRNA ARCT-154 elicited a longitudinally unique humoral profile to target (WT Spike) and to target-related Spikes, including highly diverged Omicron sublineages. This antibody response was characterized by a sustained FcγRIIIA-binding signature, which translated to an enhanced antibody-dependent natural killer cell activation (ADNKA, also referred to as antibody-dependent cellular cytotoxicity, or ADCC). These ADNKA/ADCC responses have been linked to FcγRIIIA (also known as CD16a) engagement, and ADNKA has been previously reported as predictor of protection against respiratory pathogens (29, 32, 35, 62). Future studies further characterizing how ADNKA/ADCC leveraged antibodies, either stimulated by vaccinations or provided by monoclonal antibody treatments, drive protection against respiratory pathogens are warranted.

The sustained abundance of Spike-specific antibodies elicited by ARCT-154 was notable, and while antibody profiles between the two treatment arms observed in this study showed synchronization towards one year after boost, significant changes in rates of decay and overall humoral profiles were noted between peak responses and the end of the observation period. This supports the model that a sustained production of antigen through a sa-mRNA platform influences immune response dynamics. Additionally, our results may further support a model whereby sustained antigen stimulation shifts antibody networks as a whole, similar to what is observed in hybrid immune profiles (42, 63–69). Future studies on how a sa-mRNA vaccine response compares to mRNA-vaccine- and hybrid-immune-profiles would seem warranted.

### Limitations of this Study

One limitation of this study is that it compared immune profiles of previously vaccinated participants, and we could not directly compare foundational immune profiles established by a sa-mRNA vaccine with a conventional mRNA vaccine. Separately, spacing between sampling times, particularly for later intervals, resulted in our using averaged rates of change between time points to establish vaccine dynamics. This kept us from parsing more narrowly-defined windows for peak and waned responses. It is entirely possible that peak responses to BNT162B2 boosting were before the first draw at day 29, which could have implications for our rate of change modeling and interpretations.

## Methods

### Study Design

The primary study was a double-blind, multicenter, randomized, controlled, phase 3, non-inferiority trial, conducted at 11 outpatient clinical sites in Japan (the Japan Registry for Clinical Trials, jRCT 2071220080), enrolled healthy adults aged at least 18 years who had previously been immunized with two doses of an mRNA COVID-19 vaccine (BNT162B2 [Comirnaty, Pfizer/BioNTech] or mRNA-1273 [Spikevax; Moderna]) followed by a third dose of BNT162B2 at least 3 months before enrolment. The study protocol was approved by the institutional review boards of all participating sites. The study was conducted in accordance with the ethical principles outlined in the Declaration of Helsinki and Good Clinical Practice guidelines. The results for primary and secondary endpoints were previously published (26, 28).

For this post-hoc research, we randomly selected two subsets comprising 30 participants from each vaccine group (ARCT-154 and BNT162B2, total n = 60) who were seronegative at baseline and all subsequent analysis timepoints for SARS-CoV-2 nucleocapsid protein, which is an indicator of recent COVID infection (70) and had samples available at days 1, 29, 91, 181, and 361. Samples were de-identified and processed in a blinded manner. This post-hoc testing and analysis were determined as not meeting human subjects research as determined by the Harvard IRB. All measurements were obtained from prealiquoted samples from each participant at each timepoint in each group.

### Antibody isotype and Fc-receptor binding

Antigen-specific antibody subclass/isotype levels and Fc-gamma receptor (FcγR) binding profiles were measured using a custom multiplex Luminex assay, as previously described (71–73). Carboxylated Luminex MagPlex microspheres (DiaSorin) were covalently linked to target antigens via NHS-ester linkages by the addition of EDC and Sulfo-NHS (Thermo Fisher).

Immune complex formation between serum samples diluted in 1X PBS (1:500 for Total IgG, IgG1, and IgG2; 1:250 for IgG3, IgG4, IgA1, and IgM; 1:2000 for FcγRIIA, FcγRIIB, FcγRIIIA, and FcγRIIIB) and the antigen-coupled microspheres occurred overnight at 4°C in 384-well plates, shaken at 750 rpm in 1X Assay Buffer (1X PBS, pH = 7.4, 0.1% BSA, 0.02% Tween-20). After immune complex formation, the plates were washed with 1X Assay Buffer. Ig isotypes and subclasses were detected with diluted Phycoerythrin (PE)-conjugated secondary antibodies (SouthernBiotech, see Supplementary Table 1) for 1 h at room temperature. Complexes were then washed with 1X Assay Buffer for three washes and resuspended in 1X Luminex Sheath Buffer.

FcγR-binding antibodies were detected by Avi-tagged human FcγRs (Duke Human Vaccine Institute), which were biotinylated and coupled to Streptavidin-PE (Agilent Technologies). The Strep-PE-FcγRs were incubated with the immune complexes for 1 h at room temperature in 1X Assay Buffer. Complexes were then washed with 1X Assay Buffer three times and resuspended in 1X Luminex Sheath Buffer.

All binding measurements were done using the xMAP Intelliflex (Luminex) with readouts as median fluorescence intensity (MFI) and reported in arbitrary units (AU). All samples were run in duplicate, and the means of the technical replicates were reported.

### Antibody-dependent Cellular Phagocytosis by Monocytes

Human monocytes were obtained from leukopacks (StemCell). The leukopacks were initially processed for red blood cell depletion through the EasySep RBC Depletion Reagent (StemCell). This involved adding EDTA and diluting the Leukopacks at a 1:1 ratio with EasySep Buffer (StemCell). The diluted Leukopacks then underwent a series of magnetic separations with the addition of RBC Depletion Reagent (StemCell). The cells are spun down and washed using EasySep Buffer (StemCell). Total cells were then counted using the Cell Countess, and a total of 5x10^5 cells / well were seeded onto a 96-well round-bottomed plated with R10 media (RPMI-1640 (Sigma Aldrich) supplemented with 10% fetal bovine serum (FBS) (Sigma Aldrich), 5% penicillin/streptomycin (Corning, 50 μg/mL), 5% L-glutamine (Corning, 4 mM), 5% HEPES buffer (pH 7.2) (Corning, 50 mM)).

Antigen target proteins were biotinylated using the EZ-link Sulfo-NHS-LC-LC-Biotin kit (Thermo Fisher). The biotinylated antigens were then coupled to neutravidin beads (Thermo Fisher, F8776). Next, the bead-antigen conjugates were incubated with 1:100 diluted serum overnight at 4°C to form immune complexes. The unbounded antibodies were removed using a wash buffer (1x PBS). The immune complexes were then incubated with PBMCs for 1.5 hours at 37°C 5% CO2. The cells were then stained with CD14 Pacific blue antibody cocktail at a 1:100 dilution and incubated for 30 minutes in the dark at room temperature. The stained cells were subsequently washed and fixed in 4% paraformaldehyde (PFA). The cells are washed a final time and resuspended in 1X PBS. The antigen-specific PhagoScore was determined as (% cells positive × Median Fluorescent Intensity of positive cells). Flow cytometry was performed with an iQue (IntelliCyt) instrument, and population measurements were conducted using IntelliCyt ForeCyt (v8.1). The reagents and materials used are listed in Supplementary Table 1, and the gating for ADCP is shown in Supplementary Figure 9A.

### Antibody-dependent natural killer cell (NK) activation (ADNKA)

Human natural killer (NK) cells were isolated from leukopacks (StemCell). The NK cells were enriched from the leukopacks using the EasySep Human NK Cell Isolation Kit. This is a negative selection procedure where unwanted cells are labeled with antibody complexes and magnetic rapidspheres (StemCell). The unwanted magnetically tagged cells are then separated from the desired NK cells through a series of separations using the EasySep magnets (StemCell).

Donors were anonymized. Cell concentration and viability was quantified on the Cell Countess. NK cells were resuspended in RPMI-1640 (Sigma Aldrich) supplemented with 10% fetal bovine serum (FBS) (Sigma Aldrich), 5% penicillin/streptomycin (Corning, 50 μg/mL), 5% L-glutamine (Corning, 4 mM), 5% HEPES buffer (pH 7.2) (Corning, 50 mM) in 96-well plates. The NK cells were activated overnight with the addition 1 ng/mL IL-15 (StemCell Technologies) at 37°C 5% CO2.

ELISA plates were coated with 3 μg/mL of target antigen and incubated for 2 h at 37°C. The coated plates were then washed with PBS and blocked with blocking buffer consisting of 5% bovine serum albumin (BSA) in 1X PBS, pH = 7.4, for 1 h at room temperature. The ELISA plates were washed with PBS, and 1:80 diluted serum samples were added to the plates to form immune complexes overnight at 4°C. After another wash, NK cells at a concentration of 5x10^5 in R10 media supplemented with anti-CD107a–phycoerythrin (PE)–Cy5 (BD Biosciences), brefeldin A (5 μg/mL) (Sigma-Aldrich), and GolgiStop (BD Biosciences) was added to the plates. The NK cells were incubated with immune complexes for 5 hours at 37°C 5% CO2. The incubated NK cells were subsequently washed and stained for cell surface markers with anti-CD3 Pacific Blue (BD Biosciences), anti-CD16 allophycocyanin (APC)-Cy5 (BD Biosciences), and anti-CD56 PE-Cy7 (BD Biosciences) for 20 minutes at room temperature. The washed NK cells were then fixed with PermA (Life Technologies) for another 15 minutes. The cells were washed once more and then permeabilized with PermB (Life Technologies) supplemented with anti-MIP-1β PE (BD Biosciences) and anti-IFNγ FITC for 20 minutes at room temperature.

Fluorescent intensity was measured using the iQue Cytometer (Intellicyt). NK cells were gated as CD56+/CD16+/CD3-. NK activation was evaluated as the percentage of NK cells positive for CD107a, IFNγ, or MIP-1β. All assays were performed with at least two healthy donors with one male and one female. All reagents and materials used are listed in Supplementary Table 1, and the gating strategy for ADNKA is shown in Supplementary Figure 9B.

All antibody effector function assays were quantified for statistical significance between groups (BNT162B2 and ARCT-154) using a two-sided Wilcoxon Test followed by a false discovery rate adjustment.

### Rates of change analysis

The rate of change was calculated based on the difference of the MFI within a certain period of time as shown here: 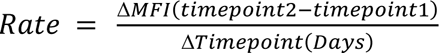. The calculation of the rate of change was paired based on the subjects, if there is no participant on certain timepoint, the data point for the time interval was not included in quantifying the group median or comparisons across groups. A two-sided Wilcoxon Test followed by a false discovery rate adjustment was used for all statistical comparisons.

### Multivariate analysis

A partial least squares discriminant analysis (PLSDA) was used to identify features separating vaccine arms post vaccination on day 1 (baseline), 29, 91, 181, and 361. Initially, all data were background subtracted, log 10 transformed, and scaled through z-scoring. The LASSO-selected features were plotted onto an LV1 loading plot to show relative contributions to the model.

Features must have been present in 80% of 100 iterations for LASSO-selection. Model performance was determined by 5-fold cross-validation, that were repeated 30 times, with comparisons to random features and permuted labels 30 times each, respectively. The validation of the LASSO-selected model was performed using the original dataset. All models (LASSO-selected, random features selected, and permuted labels selected) were quantified for accuracy from 0-1, or from 0% to 100%. The resulting score was quantified by: 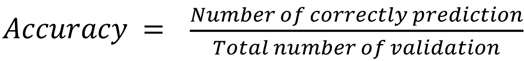. Comparisons between models to assess statistical significance favoring LASSO-selected features were done based on empirical permutation-based p-values comparing the model’s performance against a distribution of random feature or permuted label performance.

## Acknowledgements

The samples used in this research were obtained from the ARCT-154-J01 study conducted by Meiji Seika Pharma. Shinanozaka CL, Shinjuku, Japan], Hiroaki Kondo [Higashi-Shinanozaka CL, Shinjuku, Japan], Kenichi Furihata [P-One CL, Hachioji, Japan], Hidetoshi Furuie [Osaka Pharmacology Clinical Research Hospital, Osaka, Japan], Osamu Matsuoka [ToCROM, Shinjuku, Japan], Shinya Mitsui [Shin-Sapporo HP, Sapporo, Japan], Yuki Sekiguchi [LUNA, Yokohama, Japan], Shokei Kim-Mitsuyama [Medimesse Sakurajyuji, Kumamoto, Japan], Masao Kobayakawa [Fukishima Medical HP, Fukushima, Japan], and Toshio Naito [Juntendo, Bunkyo, Japan]) for their contributions to the study.

## Data Availability Statement

Raw data used for this paper has been deposited on the HSPH Systems Serology GitHub page under the accession number QW20251028 (https://github.com/HSPHSystemsSerology/QW20251028). No unique code was generated for this study. All data generated or analyzed during this study are included in this article.

## Ethics Approval

This project was determined as not human subjects research by the Harvard IRB (IRB24-1684). For the Phase III trial, all participants were enrolled through written informed consent, and in accordance with the Declaration of Helsinki. The clinical trial number for the Phase III trial is jRCT2071220080 registered on December 16, 2022.

## Author Contributions

Conceptualization: R.P.M., I.S., R.S. Methodology: K.S.L., R.B., Q.W., R.P.M.

Software: Q.W., R.P.M.

Validation: Q.W., R.P.M.

Formal Analysis: K.S.L., R.B., Q.W., R.P.M., H.J., C.V., I.S.

Investigation: All authors Resources: R.P.M., C.V., B.S.

Data Curation: K.S.L., R.B., H.M., Q.W., R.P.M.

Writing: All authors Visualization: Q.W., R.P.M.

Supervision: Q.W., R.P.M., B.S., I.S. Project Administration: H.M., S.L. Funding Acquisition: R.P.M., B.S., I.S.

