## Supplementary Information for "Sustained Humoral Activation through self-amplifying mRNA Vaccination Enhances Longitudinal Antibody Function in a Phase III Trial"

### Supplemental Information

Supplementary information includes Supplementary Table 1 and Supplementary Figures 1-9 and their corresponding captions.

### Supplementary Table 1

| REAGENT or RESOURCE | SOURCE | IDENTIFIER |
|---|---|---|
| **Antibodies** | | |
| Mouse Anti-Human IgG Fc-PE (JDC-10) | SouthernBiotech | 9040-09 |
| Mouse Anti-Human IgG1 Hinge-PE (4E3) | SouthernBiotech | 9052-09 |
| Mouse Anti-Human IgG2 Fc-PE (31-7-4) | SouthernBiotech | 9060-09 |
| Mouse Anti-Human IgG3 Hinge-PE (HP6050) | SouthernBiotech | 9210-09 |
| Mouse Anti-Human IgG4 Fc-PE (HP6025) | SouthernBiotech | 9200-09 |
| Mouse Anti-Human IgA1-PE (B3506B4) | SouthernBiotech | 9130-09 |
| Mouse Anti-Human IgA2-PE (A9604D2) | SouthernBiotech | 9140-09 |
| Mouse Anti-Human IgM-PE (SA-DA4) | SouthernBiotech | 9020-09 |
| PE-Cy™5 Mouse Anti-Human CD107a | BD Biosciences | 555802 |
| PE-Cy™7 Mouse Anti-Human CD56 (NCAM-1) | BD Biosciences | 557747 |
| APC-Cy™7 Mouse Anti-Human CD16 | BD Biosciences | 557758 |
| Pacific Blue™ Mouse Anti-Human CD3 | BD Biosciences | 558117 |
| PE Mouse Anti-Human MIP-1β | BD Biosciences | 550078 |
| FastImmune™ FITC Mouse Anti-Human IFN-γ | BD Biosciences | 340449 |
| Pacific Blue™ anti-human CD14 Antibody | BioLegend | 325616 |
| FITC IgG Goat Anti-Guinea Pig Complement C3 | MP Biomedicals | 0855385 |
| **Chemicals, Peptides, and Recombinant Proteins** | | |
| SARS-CoV-2 WT Spike Trimer | Sino Biological | 40589-V08H4 |
| SARS-CoV-2 WT Spike Trimer | Abwiz Bio | 2720-200 |
| SARS-CoV-2 KP.3 Spike Trimer | Abwiz Bio | 2804-200 |
| SARS-CoV-2 Delta Spike Trimer | Sino Biological | 4089-V08B16 |
| SARS-CoV-2 Delta Spike Trimer | Abwiz Bio | 2611-200 |
| SARS-CoV-2 BA.2 Spike Trimer | Abwiz Bio | 2460-200 |
| SARS-CoV-2 BA.5 Spike Trimer | Abwiz Bio | 2688-200 |
| SARS-CoV-2 XBB.1.5 Spike Trimer | Abwiz Bio | 2712-200 |
| Influenza A H1NA (A/Brisbane/59/2007) Hemagglutinin | Sino Biological | 11052-V08H |
| Human Cytomegalovirus Glycoprotein B | Sino Biological | 10202-V08H1 |
| Ebola Virus (subtype Zaire) Glycoprotein | Sino Biological | 40459-V08H |

| | | |
|---|---|---|
| Human soluble FcγRIIA | Duke University | Custom Order |
| Human soluble FcγRIIB | Duke University | Custom Order |
| Human soluble FcyRIIIA | Duke University | Custom Order |
| Human soluble FcyRIIIB | Duke University | Custom Order |
| Sulfo-NHS (N-hydroxysulfosuccinimide) | Thermo Fisher | A39269 |
| Pierce EDC | Thermo Fisher | A35391 |
| BirA500: BirA biotin-protein ligase standard reaction kit | Avidity LLC | BirA500 |
| Streptavidin-R-Phycoerythrin | Agilent | PJ31S |
| Brefeldin A | Sigma Aldrich | B7651 |
| Human IL-15 Recombinant Protein | Stemcell Technologies | 78031 |
| Gelatin Veronal Buffer | Sigma Aldrich | G6514 |
| Paraformaldehyde solution 4% in PBS | Santa Cruz Biotechnology | sc-281692 |
| GolgiStop™ Protein Transport Inhibitor | BD Biosciences | 554724 |
| UltraPure$^{TM}$ 0.5M EDTA, pH 8.0 | Thermo Fisher | 15575020 |
| Dimethyl Sulfoxide, Fisher BioReagents™ | Fisher Scientific | BP231 |
| RPMI-1640 Medium | Sigma Aldrich | R0883 |
| HEPES Buffer | Fisher Scientific | MT25060CI |
| Trypan Blue Solution, 0.4% (w/v) in PBS | Fisher Scientific | MT25900CI |
| L-glutamine Solution | Fisher Scientific | MT25005CI |
| Penicillin-Streptomycin Solution | Fisher Scientific | MT30001CI |
| Fetal Bovine Serum | Sigma Aldrich | F4135 |
| Bovine Serum Albumin | Sigma Aldrich | A4737 |
| Polysorbate 20 | Fisher Scientific | BP337 |
| Dulbecco's Phosphate-Buffered Salt Solution 1X | Fisher Scientific | MT21031CV |
| MES hydrate | Sigma Aldrich | M8250 |
| Sodium Hydroxide Solution | Sigma Aldrich | S8263 |
| Anhydrous sodium phosphate monobasic | Sigma Aldrich | S3139 |
| Sodium Azide | Sigma Aldrich | S2002 |
| **Biological Samples** | | |
| Guinea Pig Complement | MP Biomedicals | 08642831 |
| **Experimental Models: Cell Lines** | | |
| Human Peripheral Blood Leukopak (Tenth), Fresh | Stemcell Technologies | 200-0092 |
| Human Peripheral Blood Leukopak (Quarter), Fresh | Stemcell Technologies | 70500.2 |

| | | |
|---|---|---|
| **Critical Commercial Assays** | | |
| EZ-Link Sulfo NHS-LC-LC Biotin | Thermo Fisher | A35358 |
| Fix & Perm Cell Permeabilization Kit (Medium A) | Thermo Fisher | GAS001S100 |
| Fix & Perm Cell Permeabilization Kit (Medium B) | Thermo Fisher | GAS002S100 |
| EasySep™ RBC Depletion Reagent | Stemcell Technologies | 18170 |
| EasySep™ Human NK Cell Isolation Kit | Stemcell Technologies | 17955 |
| EasySep™ Buffer | Stemcell Technologies | 20144 |
| Zebra-Spin Desalting and Chromatography Columns | Thermo Fisher | 89882 |
| **Software and Algorithms** | | |
| iQue Forecyt 9.1 | Sartorius | 60028 |
| R Studio V 4.5.1 | R Project for Statistical Computing | RRID:SCR_000432 |
| **Other** | | |
| iQue Screener Plus | Sartorius | 11811 |
| Luminex™ xMAP INTELLIFLEX System | Thermo Fisher | APX2020 |
| MagPlex Microspheres | DiaSorin | MC12001-01 (Cataloged by region) |
| FluoSpheres™ NeutrAvidin™-Labeled Microspheres, 1.0 µm, yellow-green fluorescent (505/515), 1% solids | Thermo Fisher | f8776 |
| 384-well HydroSpeed Plate Washer | Tecan | 30190112 |
| Countess™ 3 Automated Cell Counter | Thermo Fisher | AMQAX2000 |
| Intelliflex Calibration Kit | Thermo Fisher | IFXCALK20 |
| Intelliflex Performance Verification Kit | Thermo Fisher | IFXPVERK20 |
| xMAP™ Sheath Concentrate PLUS, RUO | Thermo Fisher | 4050023 |
| iQue® Screener Plus Validation Beads | Sartorius | 91091 |
| iQue® Qsol Buffer Concentrate Solution | Sartorius | 91304 |

**Supplementary Figure 1**

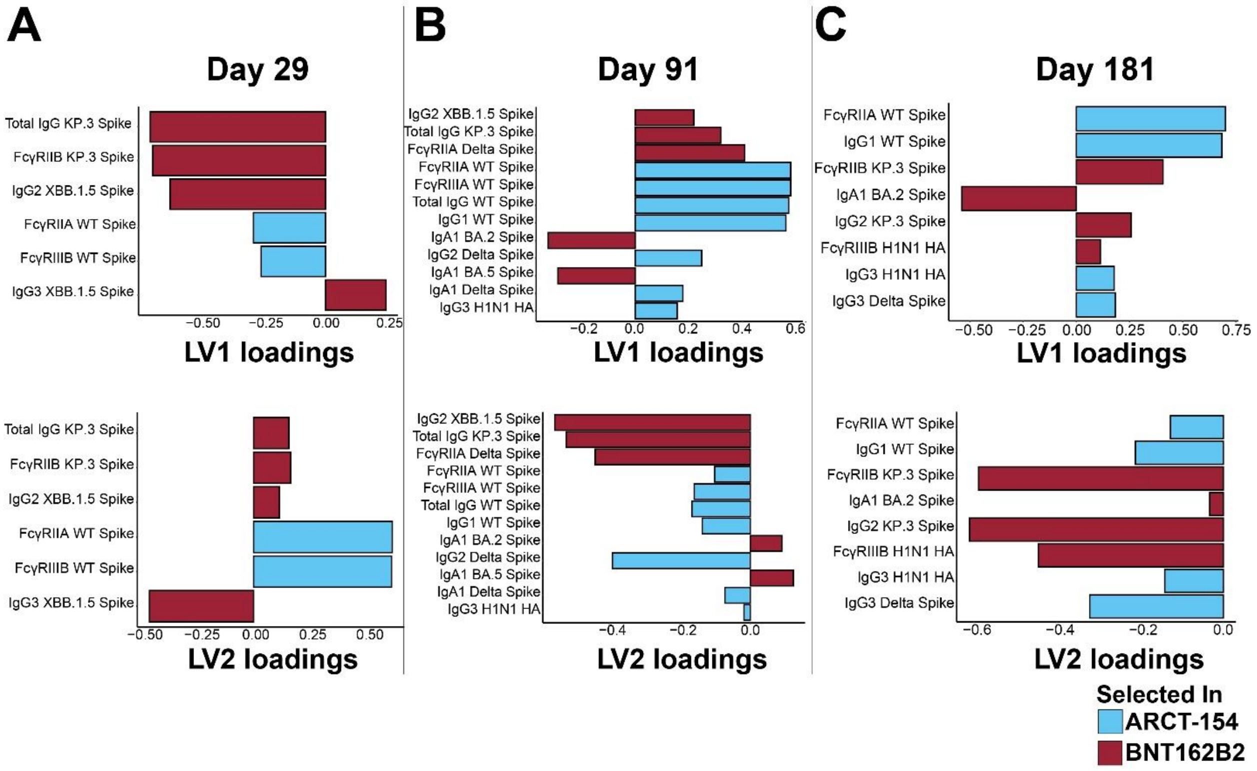


**Supplementary Figure 1. Features driving separation through a least absolute shrinkage and selection operator (LASSO)- based PLSDA.**

(A) LASSO-selected features for the PLSDA model at day 29 post-booster with BNT162B2 (red) and ARCT-154 (blue). Shown are the selected features and their scores on latent variable (LV) 1 (top) and LV2 (bottom). Shown in the bottom right corner is the color legend. No models that were statistically significant over random features and permuted labels could be made at day 1 and day 361, so no LASSO-features are shown for those time points.

(B) Same as A, but for the LASSO-selected features driving the model at day 91 post-booster.

(C) Same as A, but for the LASSO-selected features driving the model at day 181 post-booster.

**Supplementary Figure 2**

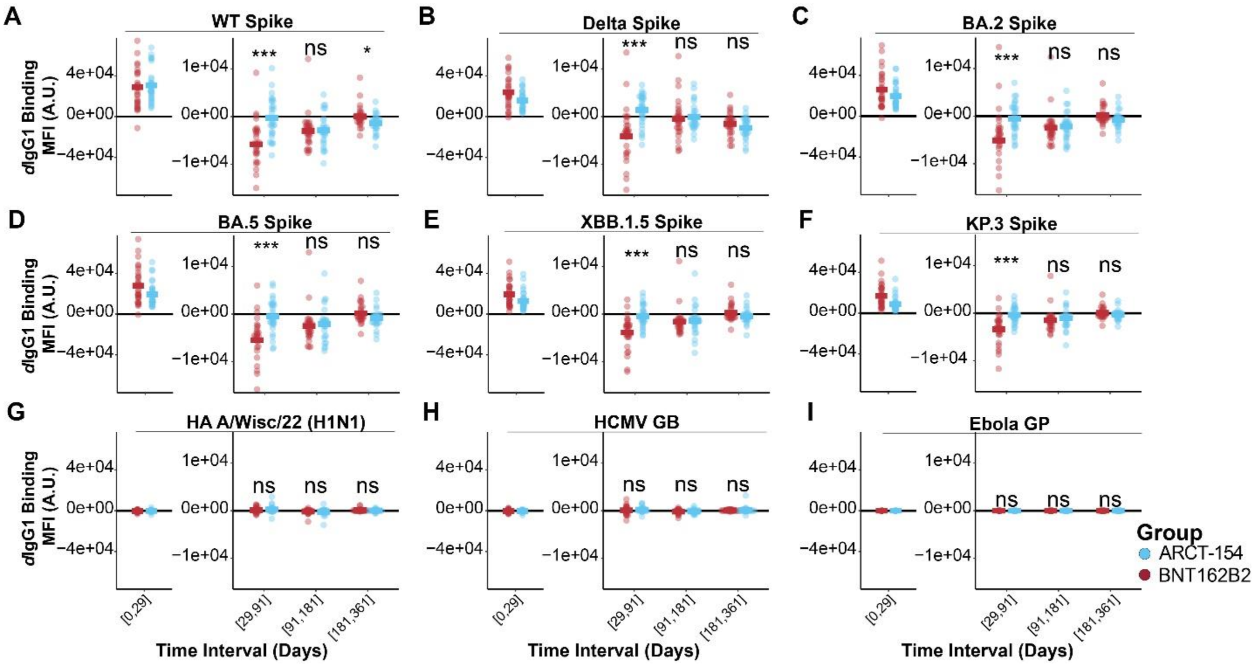


**Supplementary Figure 2. IgG1 levels to target and target-related antigens have different rates of decay based on boosting with an mRNA- or an sa-mRNA-vaccine.**

(A) Derivatives (rate of change, or *d*) were quantified for the time intervals shown on the x-axis for the two treatment arms (BNT162B2 in red, ARCT-154 in blue). Each dot represents the *d* of IgG1 to WT Spike at the time interval, and the colored horizontal bar indicates the group mean of *d* IgG1 WT Spike. A $d < 0$ indicates a negative slope, or contraction; a $d > 0$ indicates a positive slope, or expansion; and a $d =$ indicates no change in slope, or sustained response. Statistical comparisons between the two groups were done for each time interval using a Wilcoxon Test followed by a false discovery rate (FDR) adjustment. Above each time interval, n.s. indicates not statistically significant after FDR correction ($p \geq 0.05$), * indicates $p < 0.05$ after FDR correction, ** indicates $p < 0.01$ after FDR correction, and *** indicates $p < 0.001$ after FDR correction. Acute phase stimulation comparisons were not performed as this was not a primary endpoint analysis (see Methods).

(B-F) Same as A, but for variant SARS-CoV-2 Spikes shown at the top of each graph.

(G-I) Same as A, but for high-exposure control off-target antigens (HA A/Wisc/22 H1N1 and HCMV GB), and for no-exposure control antigens (Ebola GP).

### Supplementary Figure 3

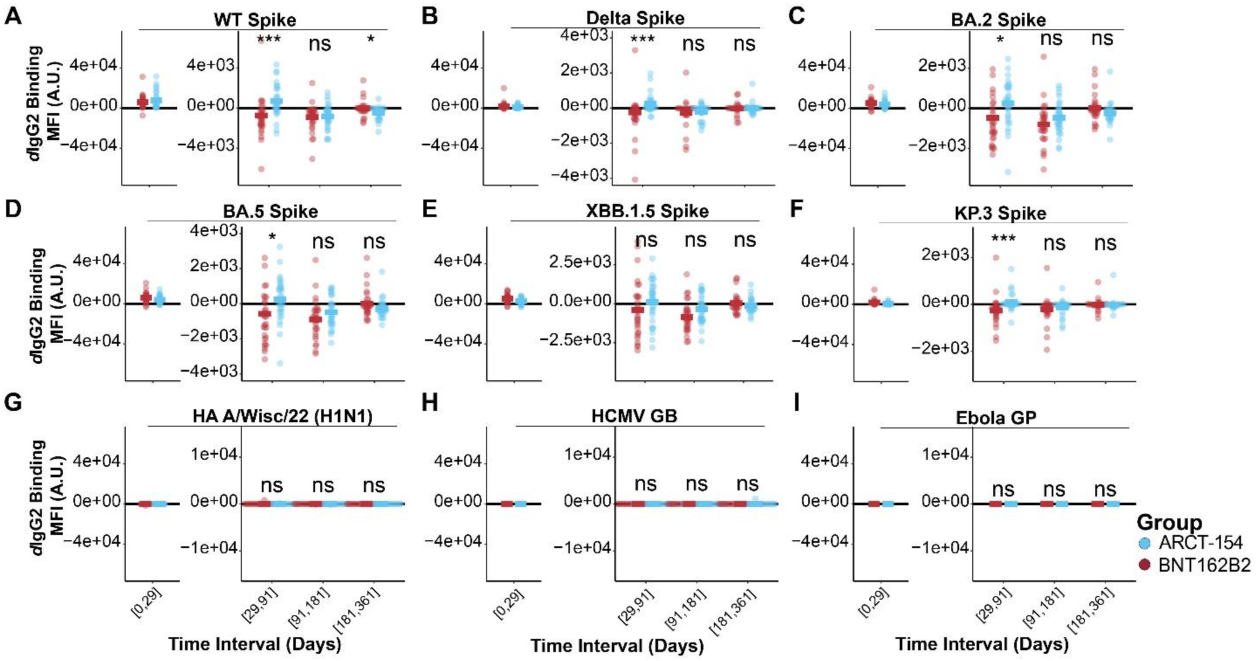


**Supplementary Figure 3. IgG2 levels to target and target-related antigens have different rates of decay based on boosting with an mRNA- or an sa-mRNA-vaccine.**

(A) Derivatives (rate of change, or *d*) were quantified for the time intervals shown on the x-axis for the two treatment arms (BNT162B2 in red, ARCT-154 in blue). Each dot represents the *d* of IgG2 to WT Spike at the time interval, and the colored horizontal bar indicates the group mean of *d* IgG2 WT Spike. A $d < 0$ indicates a negative slope, or contraction; a $d > 0$ indicates a positive slope, or expansion; and a $d =$ indicates no change in slope, or sustained response. Statistical comparisons between the two groups were done for each time interval using a Wilcoxon Test followed by a false discovery rate (FDR) adjustment. Above each time interval, n.s. indicates not statistically significant after FDR correction ($p \geq 0.05$), * indicates $p < 0.05$ after FDR correction, ** indicates $p < 0.01$ after FDR correction, and *** indicates $p < 0.001$ after FDR correction. Acute phase stimulation comparisons were not performed as this was not a primary endpoint analysis (see Methods).

(B-F) Same as A, but for variant SARS-CoV-2 Spikes shown at the top of each graph.

(G-I) Same as A, but for high-exposure control off-target antigens (HA A/Wisc/22 H1N1 and HCMV GB), and for no-exposure control antigens (Ebola GP).

### Supplementary Figure 4

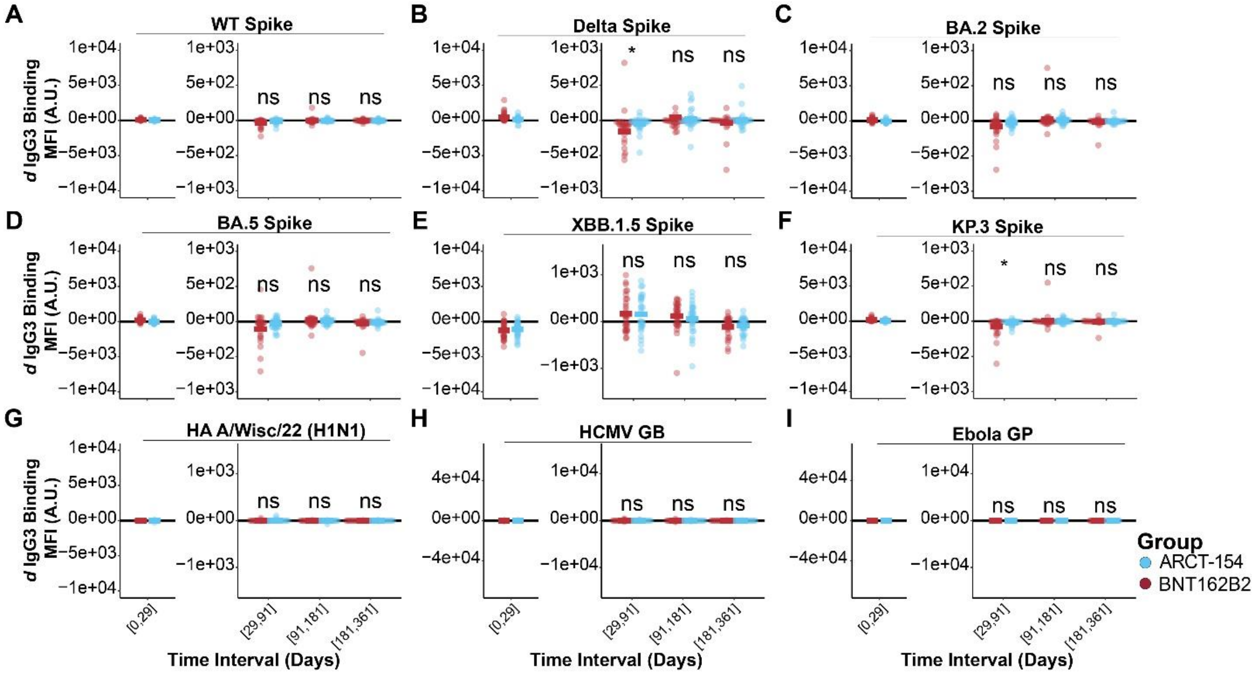


**Supplementary Figure 4. IgG3 levels to target and target-related antigens have different rates of decay based on boosting with an mRNA- or an sa-mRNA-vaccine.**

(A) Derivatives (rate of change, or *d*) were quantified for the time intervals shown on the x-axis for the two treatment arms (BNT162B2 in red, ARCT-154 in blue). Each dot represents the *d* of IgG3 to WT Spike at the time interval, and the colored horizontal bar indicates the group mean of *d* IgG3 WT Spike. A $d < 0$ indicates a negative slope, or contraction; a $d > 0$ indicates a positive slope, or expansion; and a *d* = indicates no change in slope, or sustained response. Statistical comparisons between the two groups were done for each time interval using a Wilcoxon Test followed by a false discovery rate (FDR) adjustment. Above each time interval, n.s. indicates not statistically significant after FDR correction ($p \geq 0.05$), * indicates $p < 0.05$ after FDR correction, ** indicates $p < 0.01$ after FDR correction, and *** indicates $p < 0.001$ after FDR correction. Acute phase stimulation comparisons were not performed as this was not a primary endpoint analysis (see Methods).

(B-F) Same as A, but for variant SARS-CoV-2 Spikes shown at the top of each graph.

(G-I) Same as A, but for high-exposure control off-target antigens (HA A/Wisc/22 H1N1 and HCMV GB), and for no-exposure control antigens (Ebola GP).

### Supplementary Figure 5

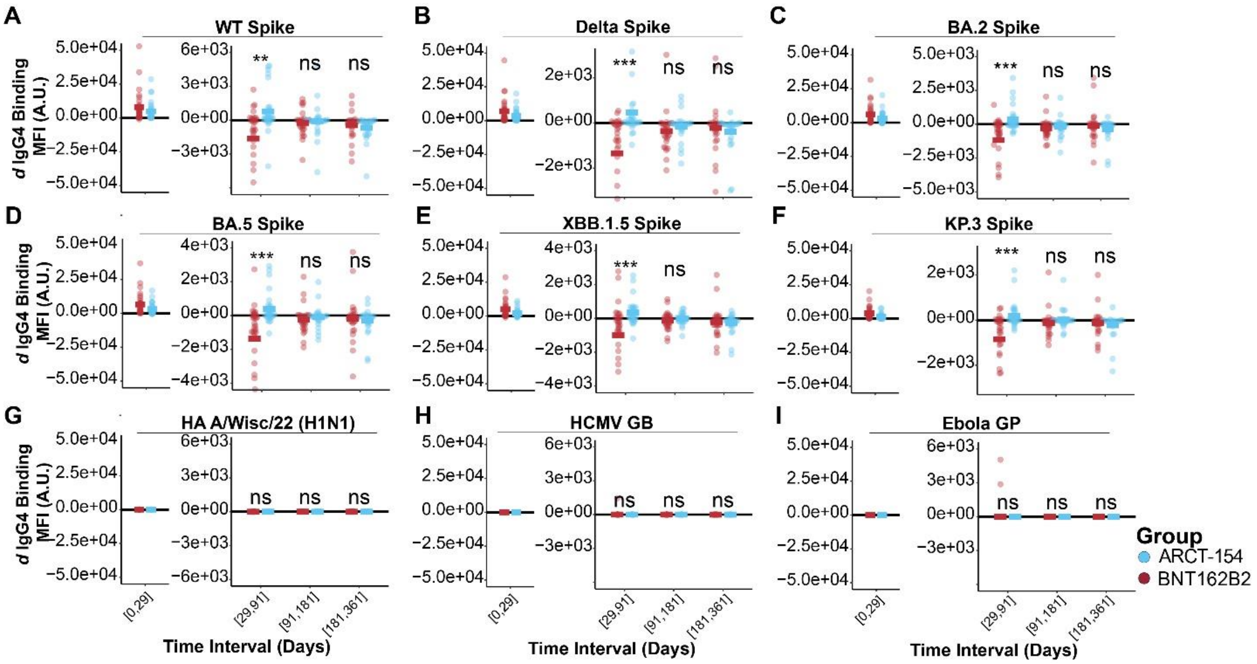


**Supplementary Figure 5. IgG4 levels to target and target-related antigens have different rates of decay based on boosting with an mRNA- or an sa-mRNA-vaccine.**

(A) Derivatives (rate of change, or *d*) were quantified for the time intervals shown on the x-axis for the two treatment arms (BNT162B2 in red, ARCT-154 in blue). Each dot represents the *d* of IgG4 to WT Spike at the time interval, and the colored horizontal bar indicates the group mean of *d* IgG4 WT Spike. A $d < 0$ indicates a negative slope, or contraction; a $d > 0$ indicates a positive slope, or expansion; and a $d =$ indicates no change in slope, or sustained response. Statistical comparisons between the two groups were done for each time interval using a Wilcoxon Test followed by a false discovery rate (FDR) adjustment. Above each time interval, n.s. indicates not statistically significant after FDR correction ($p \geq 0.05$), * indicates $p < 0.05$ after FDR correction, ** indicates $p < 0.01$ after FDR correction, and *** indicates $p < 0.001$ after FDR correction. Acute phase stimulation comparisons were not performed as this was not a primary endpoint analysis (see Methods).

(B-F) Same as A, but for variant SARS-CoV-2 Spikes shown at the top of each graph.

(G-I) Same as A, but for high-exposure control off-target antigens (HA A/Wisc/22 H1N1 and HCMV GB), and for no-exposure control antigens (Ebola GP).

### Supplementary Figure 6

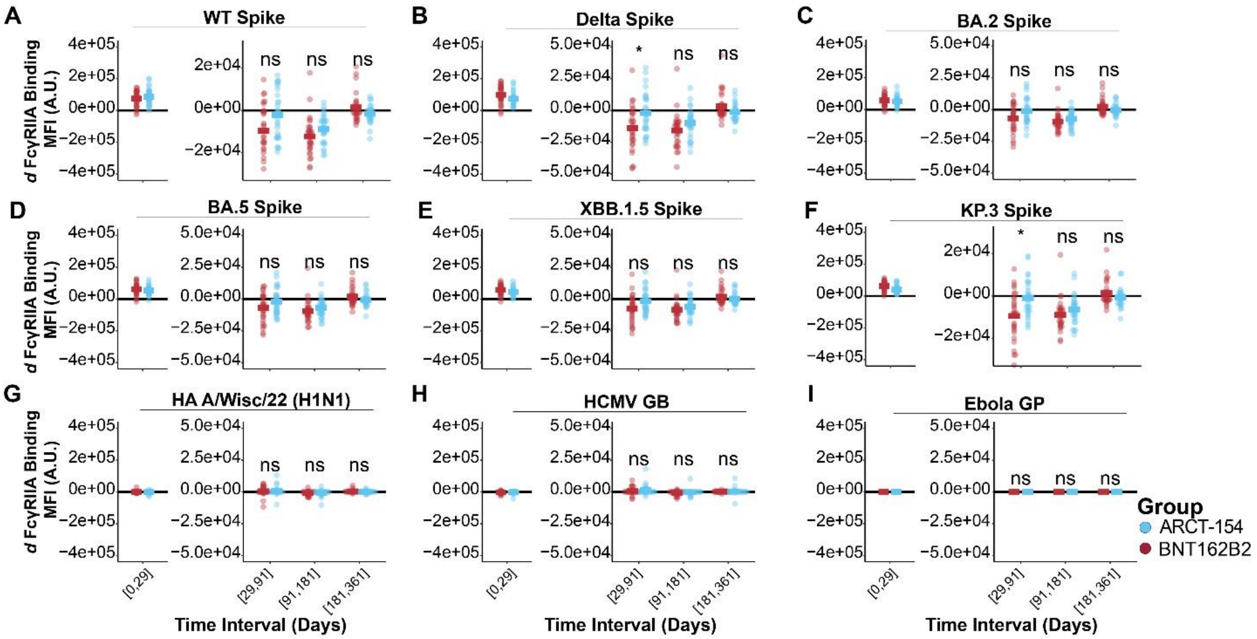


**Supplementary Figure 6. FcγRIIA-binding antibody levels to target and target-related antigens have different rates of decay based on boosting with an mRNA- or an sa-mRNA-vaccine.**

(A) Derivatives (rate of change, or *d*) were quantified for the time intervals shown on the x-axis for the two treatment arms (BNT162B2 in red, ARCT-154 in blue). Each dot represents the *d* of FcγRIIA-binding antibodies to WT Spike at the time interval, and the colored horizontal bar indicates the group mean of *d* FcγRIIA-binding antibodies to WT Spike. A $d < 0$ indicates a negative slope, or contraction; a $d > 0$ indicates a positive slope, or expansion; and a *d* = indicates no change in slope, or sustained response. Statistical comparisons between the two groups were done for each time interval using a Wilcoxon Test followed by a false discovery rate (FDR) adjustment. Above each time interval, n.s. indicates not statistically significant after FDR correction ($p \geq 0.05$), * indicates $p < 0.05$ after FDR correction, ** indicates $p < 0.01$ after FDR correction, and *** indicates $p < 0.001$ after FDR correction. Acute phase stimulation comparisons were not performed as this was not a primary endpoint analysis (see Methods).

(B-F) Same as A, but for variant SARS-CoV-2 Spikes shown at the top of each graph.

(G-I) Same as A, but for high-exposure control off-target antigens (HA A/Wisc/22 H1N1 and HCMV GB), and for no-exposure control antigens (Ebola GP).

**Supplementary Figure 7**

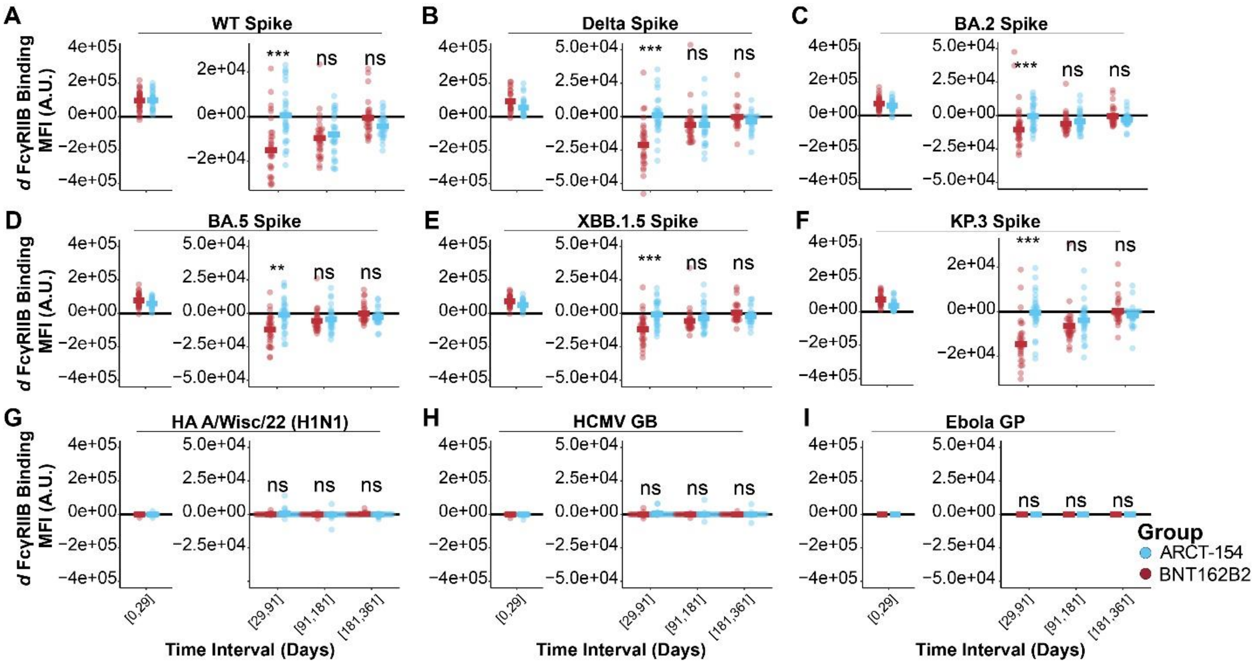


**Supplementary Figure 7. FcγRIIB-binding antibody levels to target and target-related antigens have different rates of decay based on boosting with an mRNA- or an sa-mRNA-vaccine.**

(A) Derivatives (rate of change, or *d*) were quantified for the time intervals shown on the x-axis for the two treatment arms (BNT162B2 in red, ARCT-154 in blue). Each dot represents the *d* of FcγRIIB-binding antibodies to WT Spike at the time interval, and the colored horizontal bar indicates the group mean of *d* FcγRIIB-binding antibodies to WT Spike. A $d < 0$ indicates a negative slope, or contraction; a $d > 0$ indicates a positive slope, or expansion; and a *d* = indicates no change in slope, or sustained response. Statistical comparisons between the two groups were done for each time interval using a Wilcoxon Test followed by a false discovery rate (FDR) adjustment. Above each time interval, n.s. indicates not statistically significant after FDR correction ($p \geq 0.05$), * indicates $p < 0.05$ after FDR correction, ** indicates $p < 0.01$ after FDR correction, and *** indicates $p < 0.001$ after FDR correction. Acute phase stimulation comparisons were not performed as this was not a primary endpoint analysis (see Methods).

(B-F) Same as A, but for variant SARS-CoV-2 Spikes shown at the top of each graph.

(G-I) Same as A, but for high-exposure control off-target antigens (HA A/Wisc/22 H1N1 and HCMV GB), and for no-exposure control antigens (Ebola GP).

**Supplementary Figure 8**

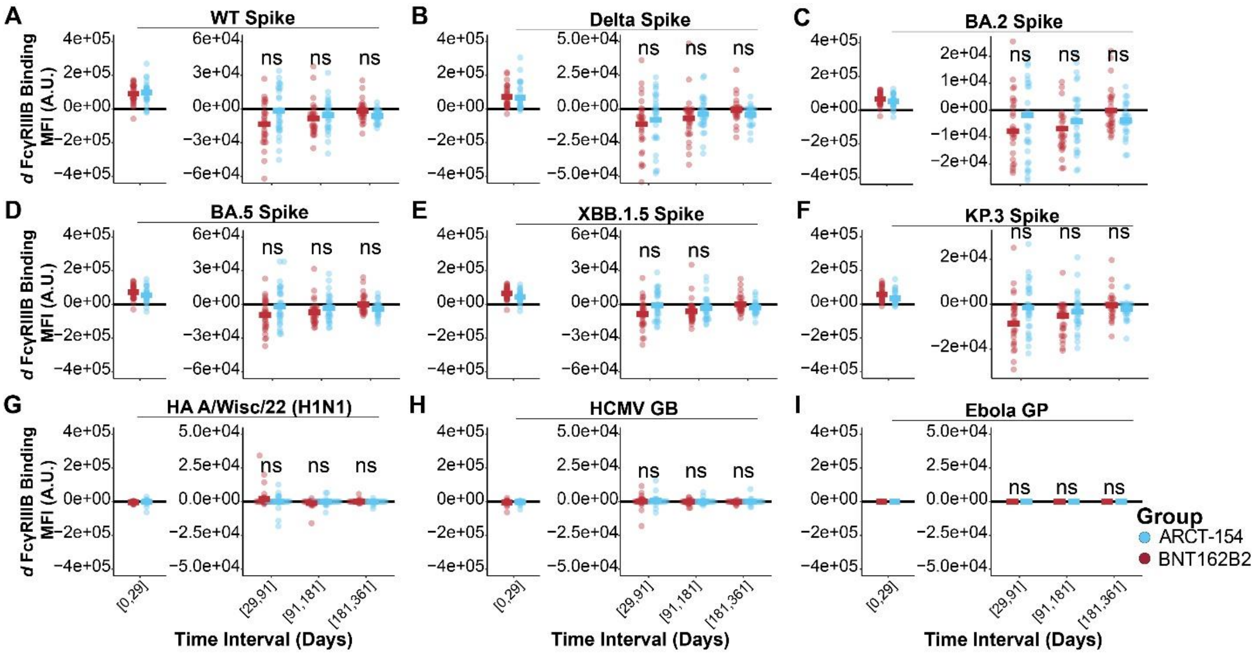


**Supplementary Figure 8. FcγRIIIB-binding antibody levels to target and target-related antigens have different rates of decay based on boosting with an mRNA- or an sa-mRNA-vaccine.**

(A) Derivatives (rate of change, or *d*) were quantified for the time intervals shown on the x-axis for the two treatment arms (BNT162B2 in red, ARCT-154 in blue). Each dot represents the *d* of FcγRIIIB-binding antibodies to WT Spike at the time interval, and the colored horizontal bar indicates the group mean of *d* FcγRIIIB-binding antibodies to WT Spike. A $d < 0$ indicates a negative slope, or contraction; a $d > 0$ indicates a positive slope, or expansion; and a $d =$ indicates no change in slope, or sustained response. Statistical comparisons between the two groups were done for each time interval using a Wilcoxon Test followed by a false discovery rate (FDR) adjustment. Above each time interval, n.s. indicates not statistically significant after FDR correction ($p \geq 0.05$), * indicates $p < 0.05$ after FDR correction, ** indicates $p < 0.01$ after FDR correction, and *** indicates $p < 0.001$ after FDR correction. Acute phase stimulation comparisons were not performed as this was not a primary endpoint analysis (see Methods).

(B-F) Same as A, but for variant SARS-CoV-2 Spikes shown at the top of each graph.

(G-I) Same as A, but for high-exposure control off-target antigens (HA A/Wisc/22 H1N1 and HCMV GB), and for no-exposure control antigens (Ebola GP).

**Supplementary Figure 9**

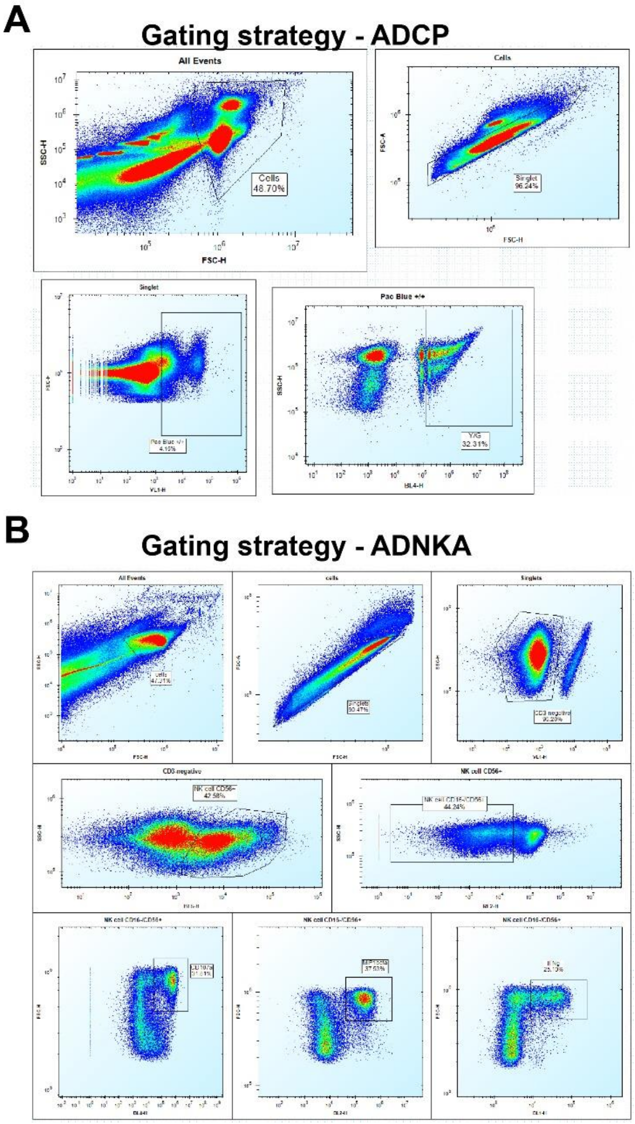


**Supplementary Figure 9. Flow cytometry gating for antibody effector assays.**

(A) Gating strategy for ADCP from primary-derived human leukopacks. Singlets were gated, and CD14+ cells were further gated to quantify monocyte responses.

(B) Gating strategy for ADNKA from primary-derived human leukopacks. Singlets were gated, and CD56+/CD3- NK cells were further gated to quantify NK responses.
